# Visual cues determine hippocampal directional selectivity

**DOI:** 10.1101/017210

**Authors:** Lavanya Acharya, Zahra M. Aghajan, Cliff Vuong, Jason J. Moore, Mayank R. Mehta

## Abstract

Both spatial and directional information are necessary for navigation. Rodent hippocampal neurons show spatial selectivity in all environments^1^, but directional tuning only on linear paths^2–8^.The sensory mechanisms underlying directionality are unknown, though vestibular and visual cues are thought to be crucial. However, hippocampal neurons are thought to show no angular modulation during two-dimensional random foraging despite the presence of vestibular and visual cues^6,7^. Additionally, specific aspects of visual cues have not been directly linked to hippocampal responses in rodents. To resolve these issues we manipulated vestibular and visual cues in a series of experiments. We first measured hippocampal activity during random foraging in real world (RW) where we found that neurons’ firing exhibited significant modulation by head-direction. In fact, the fraction of modulated neurons was comparable to that in the head-direction system^9^. These findings are contrary to commonly held beliefs about hippocampal directionality^6,7^. To isolate the contribution of visual cues we measured neural responses in a visually similar virtual reality (VR) where the range of vestibular inputs is minimized^5,10,11^. Significant directional modulation was not only found in VR, but it was comparable to that in RW. Several additional experiments revealed that changes in the angular information contained in the visual cues induced corresponding changes in hippocampal head-directional modulation. Remarkably, for head-directionally modulated neurons, the ensemble activity was biased towards the sole visual cue. These results demonstrate that robust vestibular cues are not required for hippocampal directional selectivity, while visual cues are not only sufficient but also play a causal role in driving hippocampal responses.

## Introduction

Hippocampal spatial selectivity has been well established and the underlying mechanisms extensively studied^1,12^. However, both the existence and mechanisms of hippocampal directional selectivity are debated. During random foraging in two-dimensional arenas, barring a few conflicting reports^3,6,13^, the consensus is that rodent hippocampal neurons do not show significant angular selectivity^6,7^. In contrast, hippocampal neurons exhibit strong directional selectivity on linear paths^3–5,8^. The reason for this disparity and the sensory mechanisms of directionality are unclear, although visual and vestibular cues have been proposed as likely candidates^3,5,14^. In addition, internal mechanisms also contribute to hippocampal activity^11,15–17^.

Visual cues strongly influence the spatial firing properties of hippocampal neurons ^2,18^. Further, comparable levels of directionality exist on linear tracks in RW and in VR^5^ where the range of vestibular inputs is minimal, suggesting that visual cues also influence directionality in one dimension. In addition, selectivity to the visual cue towards which the animal’s head is facing, referred to as spatial-view, has been reported in humans^19^, primates^20,21^ and bats^22^. However, response to specific features of visual cues has not been observed in rodents, leading to the notion that in these animals visual cues merely provide a context for hippocampal activity.

In parallel, vestibular inputs are crucial to the head-direction system, which is thought to provide directional information to the hippocampus. Consistently, vestibular lesions disrupt hippocampal spatial selectivity^23^, although lesions in the head-direction system do not^24^. However, if instantaneous vestibular cues were contributing to hippocampal directionality, there should be greater directionality in two-dimensional RW tasks, where the range is higher compared to one-dimensional RW tasks, but the opposite is true. Some studies have attributed directionality in two dimensions to vestibular-derived self-motion information^3,14,25^, but no study, to our knowledge, has directly measured hippocampal head-directional modulation when vestibular-based signals are impaired.

Thus, the mechanisms governing hippocampal directional activity in rodents are unclear. We hypothesize that visual cues directly influence the activity of hippocampal neurons to generate angular tuning whereas vestibular cues are not required for directionality.

## Results

To test these hypotheses we did a series of experiments and analyses. We first quantified hippocampal spatial and head-directional modulation from 1066 active (defined as cells with minimum mean firing rate of 0.2Hz and with at least a 100 spikes) dorsal CA1 pyramidal neurons (which were part of a previous study of hippocampal spatial selectivity^11^). Rats randomly foraged for rewards on a two-dimensional platform in a RW environment which had rich distal visual cues and will henceforth be referred to as RW_rich_ (Fig. 1a).

**Figure 1:**
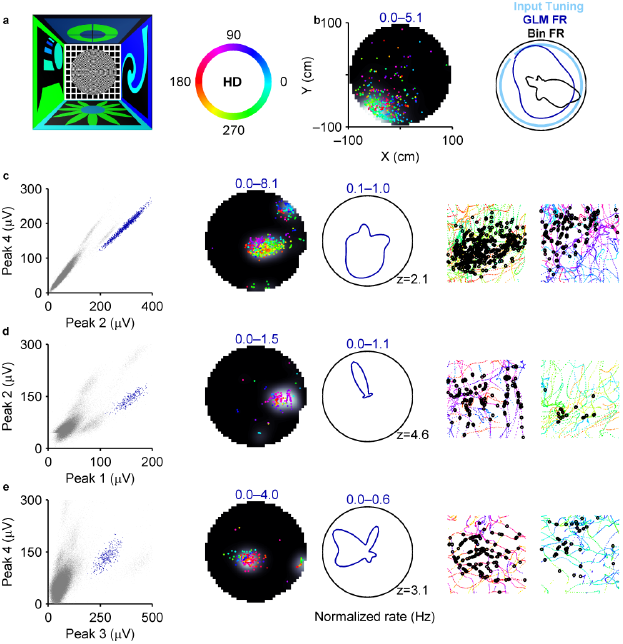
Presence of head-directional modulation in hippocampal pyramidal neurons in RW_rich_. a, Left) A top-view schematic depicting a 300cm×300cm room with four different visual cues on the walls and an elevated 100cm radius platform at the center. Right) A color wheel representing the mapping between head-directions and colors. **b,** Left) Spatial firing rate of a surrogate neuron (grey scale range indicated by numbers; lighter shades correspond to higher values, here and throughout all figures) overlaid with the position of the rat when spikes occurred (colored dots). Each color represents a distinct head-direction as shown in (**a**). The surrogate neuron’s activity was constructed to have significant spatial selectivity but no angular selectivity. Right) Angular rate map of the same surrogate neuron estimated using the binning (black) and GLM (blue) methods, along with the uniform input tuning (light blue). The GLM method provided an accurate estimate of the input, but the binning method overestimated the angular tuning due to behavioral bias (Extended Data Fig. 1). **c,** Left) All unclustered (grey dots) and clustered spike amplitudes from an isolated neuron (blue dots) on two different projections of a tetrode in RW_rich_. Center) Spatial and angular rate maps of a cell (same convention as in (**b**)). Numbers in color indicate range, here and throughout unless noted otherwise. The number at the bottom right corner of the polar plot is the z-scored sparsity of the angular rate map). Right) Rat’s color-coded trajectory and his position at the time of spikes (black circles) for movement in the direction of maximal (left) and minimal (right) firing respectively. **d,e,** Same as (**c**) for two other cells in RW_rich_. All rate maps were computed using the GLM method here and throughout unless otherwise noted.

A common technique for quantifying head-directional modulation is to divide the number of spikes in each direction bin by the total time spent in that bin (Fig. 1b)^9^. However, when neurons have spatially tuned responses, as is the case for hippocampal neurons in RW, this method provides incorrect estimates of angular tuning^6^. For example, for a neuron with a place field at the edge of the maze, this method would yield artificially large head-directional tuning due to non-uniform sampling of head angles within the place field (Fig. 1b, Extended Data Fig. 1)^6^. Various methods have been developed to overcome this confound^3,25,26^. Here, we adopted the well-established generalized linear model (GLM) approach (see Methods)^15,27–29^ which has several advantages. First, it provides an unbiased estimate of the simultaneous and independent contribution of spatial and head-directional modulation. Second, unlike other methods, head-directional modulation obtained with the GLM method is uninfluenced by behavioral biases within the place field as verified using surrogate data with predetermined levels of spatial and angular modulation (see Methods, Extended Data Fig. 1i). Finally, this method provides an estimate of the fine structure of the respective tuning curves.

This method revealed a surprising finding: many neurons exhibited clear modulation by the rat’s head-direction in RW_rich_(Fig. 1c,d,e, Extended Data Fig. 2a, see Methods). Some neurons fired maximally for only one head-direction and minimally elsewhere (Fig. 1c,d), while others showed a multimodal response (Fig. 1e). The statistical significance of head-directional modulation was quantified by the sparsity of angular rate maps z-scored with respect to control data (see Methods). Angular sparsity thus defined was significantly (*p*<0.05) greater than chance (>2z) for 26% of neurons in RW_rich_. In fact, the fraction of hippocampal neurons with significant angular tuning was comparable to that in many parts of the head-direction system, although the width of the angular tuning curves (full width at half maximum 103.10±3.50°) was wider^9,30^.

**Figure 2:**
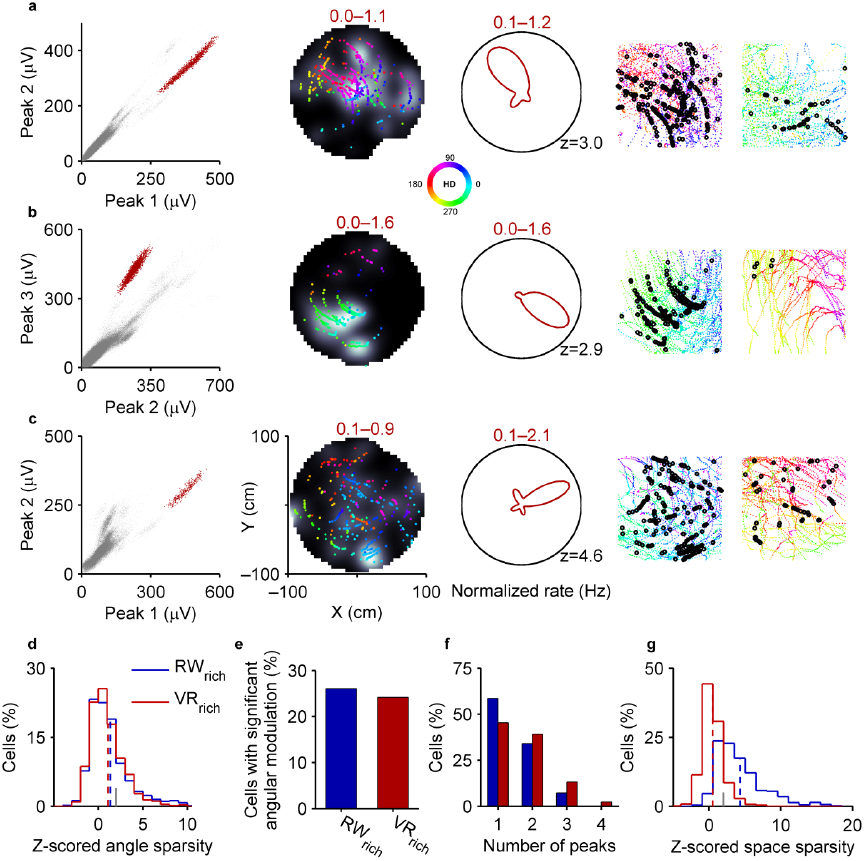
Similar levels of significant head-directional modulation in RW_rich_ **and** VR_rich_. a–c. Three well-isolated neurons showing significant head-directional modulation in VR_rich_ (same conventions as in Fig. 1). **d,** Z-scored angular sparsity in VR_rich_ (1.09±0.08, n=719, red) was similar (p=0.2) to that in RW_rich_ (1.35±0.08, n=1066, blue). Mean values are indicated with dashed vertical lines. Grey tick mark indicates significance threshold (z=2). **e,** 26% (24%) of cells in RW_rich_ (VR_rich_) showed significant head-directional modulation (see Methods). **f,** Head-directionally modulated neurons in VR_rich_ were significantly more multimodal (1.72±0.06 peaks, p=1.5×10^−3^) than RW_rich_ cells (1.49±0.04 peaks). **g,** In contrast to similar z-scored angular sparsity, z-scored spatial sparsity was significantly reduced in VR_rich_ (0.51±0.06, p=1.2×10^−174^) compared to RW_rich_ (4.32±0.11). Throughout the figure legends values are reported as mean±s.e.m and the statistical significance for comparisons was computed using Wilcoxon rank-sum test unless otherwise stated.

This raises an important question: which sensory inputs could generate the head-directional modulation in our data? Two likely candidates are the visual and vestibular modalities. To dissociate the two, we measured the activity of 719 active dorsal hippocampal CA1 pyramidal neurons^11^ during the same random foraging task in a two-dimensional VR environment (VR_rich_). Here, the distal visual cues were identical to those in RW_rich_, but the range of vestibular cues was minimized due to body fixation. Despite impaired spatial selectivity^11^, many neurons showed clear modulation by the direction of the rat’s head with respect to the distal visual cues, which will be henceforth referred to as “head-direction"(Fig.2a,b,c, Extended Data Fig. 2b).

Curiously, across the ensemble of neurons there was no substantial difference in head-directional modulation between the two worlds as quantified by z-scored angular sparsity (Fig. 2d). Further, the fraction of neurons showing significant directional modulation in VR_rich_ (24%) was similar to that in RW_rich_ (Fig. 2e). Neurons in VR_rich_ also had multimodal responses like in RW_rich_, unlike neurons in the head-direction network which have unimodal responses^31^. The multimodality was greater in VR_rich_ than RW_rich_ (Fig. 2f) which could account for slightly lower z-scored mean vector length in the former (Extended Data Fig. 3). This observation motivates the use of z-scored sparsity as a measure for angular selectivity. Additionally, the width of the angular tuning curves in VR_rich_ (86.61±4.19°) was significantly (16%, p=4.3×10^−4^) sharper than in RW_rich_.

**Figure 3:**
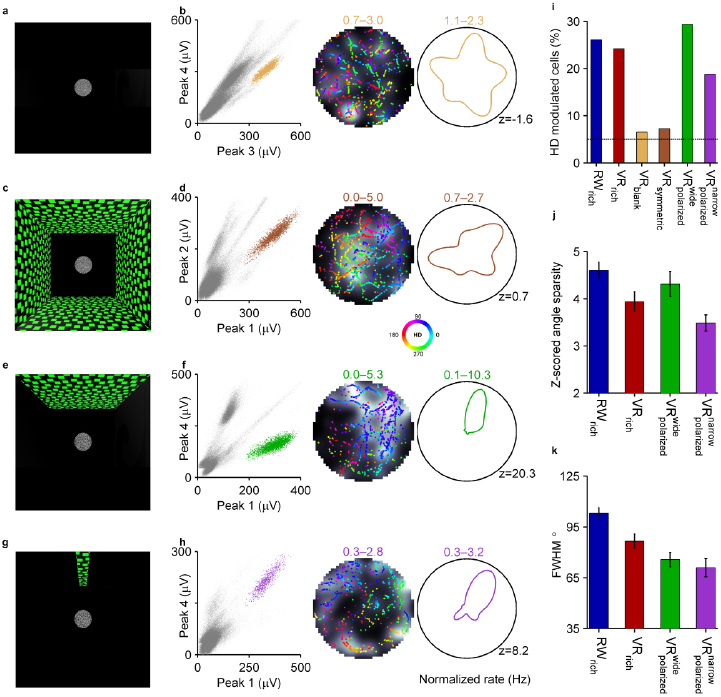
Direct influence of visual cues on the degree of directional modulation of neurons. **a**, Top-down schematic of VR task with a 100cm radius circular platform with no distal visual cues (VR_blank_). **b,** Left) Spikes from an isolated neuron (mustard dots) in VR_blank_ (same convention as in Fig. 1). Center, right) Spatial and angular firing rate of this neuron (same conventions as in Fig. 1). **c,** Top-down schematic of VR task with symmetric cues located 450cm away from the center of the circular platform (VR_symmetric_). **d,** Same as in (**b**) but in VR_symmetric_. Note that the neurons shown in both (**b, d**) do not exhibit any head-directional modulation (z<2). **e,** Top-down schematic of VR task with a single wide polarizing visual cue 450cm away from the center (90° visual angle) of the platform 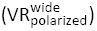. **f,** Same as (**b, d**) but in 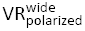 Top-down schematic of VR task with a narrow polarizing cue 450cm away from the center (11° visual angle) of the circular table 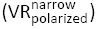. **h,** Same as (**b, d, f**) but in 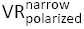. Note that the neurons shown in both (**f, h**) exhibit strong head-directional modulation (z>2). **i,** The percentages of cells with significant head-directional modulation was 26% in RW_rich_ (278 out of 1066 cells), 24% in VR_rich_ (174 of 719 cells), 6% in VR_blank_ (14 of 230 cells), 7% in VR_symmetric_ (30 of 426 cells), 30% in 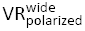 (89 of 300 cells) and 19% in 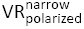 (65 of 341 cells). The black horizontal line indicates the chance level of 5%. The distribution of z-scored angular sparsity was significantly different from zero in RW_rich_ (p=3.0×10^−59^ Wilcoxon signed-rank test here and throughout (**i**)), VR_rich_ (p=3.0×10^−37^), 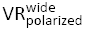 and 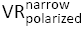 but not in VR_blank_(p=0.7) or VR_symmetric_ (p=0.2). **j,** Z-scored sparsity of the angular rate maps for cells with significant head-directional modulation in different tasks was 4.61±0.17 in RW_rich_, 3.94±0.21 in VR_rich_, 4.31±0.26 in 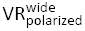 and 3.49±0.17 in 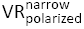 (error bars indicate s.e.m). Out of six possible pair-wise comparisons between the distributions, the only significant differences were that z-scored angular sparsity in RW_rich_ was slightly greater than VR_rich_ (p=4.1×10^−3^) and 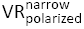. **k,** Full width at half max (FWHM) of the angular rate maps for head-directionally modulated neurons in different conditions was as follows: RW_rich_ (103.10±3.50°), VR_rich_ (86.61±4.19°), 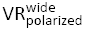 and 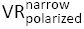. The tuning curves in RW_rich_ were significantly wider than all other VR conditions (p=4.3×10^−4^ versus VR_rich_, p=4.9×10^−5^ versus 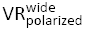 and p=6.8×10^−6^ versus 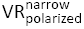). Within VR conditions, the only significant difference was polarized polarized polarized observed between VR_rich_ and 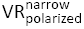 with the latter having significantly narrower tuning curves (p=0.03). Values are reported as mean±s.e.m and the p-values are obtained by Wilcoxon rank-sum test unless noted otherwise.

We then quantified the spatial modulation of neural responses in both RW_rich_ and VR_rich_ using the rate maps obtained from the GLM method. We found significant spatial selectivity in RW_rich_ but not in VR_rich_ (Fig. 2g), consistent with previous results obtained using the binning method^11^. Although head-directional modulation was comparable between the two worlds, spatial modulation was not, which suggests a decoupling of the mechanisms of spatial and directional tuning. Consistently, the presence or absence of head-directional modulation had no effect on the percentage of spatially modulated neurons in both RW_rich_ and VR_rich_(Extended Data Fig. 4a).

**Figure 4:**
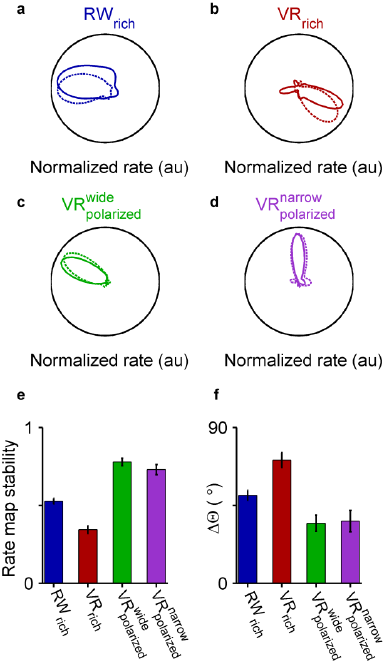
Stability of angular rate maps for head-directionally modulated neurons. a–d, Four directionally tuned cells with stable angular firing in the first half (solid colored lines) and second half (dashed colored lines) of the recording session. The peak rates are normalized for ease of comparison. **e,** Stability of the head-directional modulation (computed as the pairwise correlation between the angular rate maps in the first and second halves) in RW_rich_ (0.53±0.02, n=278) was significantly greater than VR_rich_ (0.34±0.02, n=174, p=2.2×10^−9^, Wilcoxon rank-sum test here and throughout figure legend) but significantly smaller than 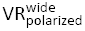 (0.78±0.02, n=89, p=1.3×10^−16^) and 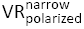(0.73±0.03, n=65, p=3.2×10^−9^). Angular rate map stability was not significantly different between 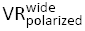 and 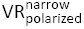(p=0.2). **f,** As an alternate measure of stability, we computed the absolute value of the circular distance between the preferred directions (defined as the direction of peak firing) in the two session halves. This method also resulted in a similar trend with 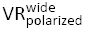 (34.59±4.75°) and 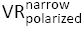 (35.89±6.19°) showing identical levels of drift of the preferred directions (p=0.5), both smaller than the amount of angular drift in RW_rich_ (50.78±2.94°, p=4.3×10^−5^ and p=1.1×10^−4^ respectively) and VR_rich_ (71.16±4.23°, p=1.4×10^−8^ and p=4.7×10^−7^ respectively).

Interestingly, directionally tuned neurons had greater mean firing rates than the untuned neurons in VR_rich_ (Extended Data Fig. 4b). This difference was not present in RW_rich_, which might be due to the presence of multisensory spatially informative cues^11^.

These results show that rodent hippocampal neurons in RW indeed show significant head-directional modulation during two-dimensional random foraging, contrary to previous reports. In addition, the observation that head-directional modulation remained intact in VR—where the range of vestibularcues is minimized—suggests that vestibular cues are not required for hippocampal head-directional modulation.

What other mechanism could generate angular modulation? Either it is internally generated^15–17^ or driven by specific visual cues^20,21^. To disambiguate these possibilities, we generated a virtual world where distal visual cues were entirely eliminated (VR_blank_)(Fig. 3a, see Methods). The circular platform in the virtual environment, which was the only visual cue present, provided optic flow information but had no spatial or angular information. The rats’ behavior in VR_blank_ was comparable to that in VR_rich_ with visually distinct walls (Extended Data Fig. 5a). Hippocampal neurons did not show clear head-directional modulation in this case (Fig. 3b,i, Extended Data Fig. 5b) and the distribution of z-scored angular sparsity was not significantly different from zero (p=0.7, Wilcoxon signed-rank test).

**Figure 5:**
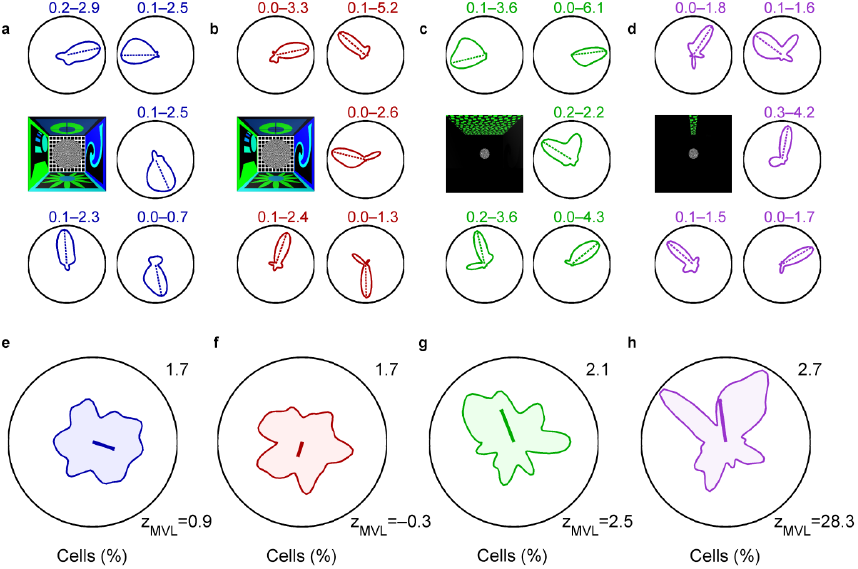
Bias of the neural ensemble by polarizing visual cues. a, Five example cells in RW_rich_ with significant directional tuning. The numbers indicate firing rate range. The dashed line corresponds to the preferred direction, i.e. the direction of maximum firing. The schematic in the middle left indicates the experimental condition. **b–d,** Same as in (**a**) but in VR_rich_, 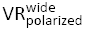 and 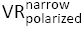 conditions respectively. **e,** Distribution of preferred direction of neurons in RW_rich_(341.71±4.66°, n=278, circular mean±circular s.e.m). The number on the top right indicates the maximum value of the distribution. The thick blue line originating at the center of the polar plot represents both the direction (341.71°) and the magnitude (0.08) of the mean vector length of the preferred directions of the population (scaled by a factor of 5 for clarity). The mean vector length was not significantly different from chance (z=0.95). **f,** Same as in **(e)** but for VR_rich_. The distribution of preferred directions of neurons in VR_rich_ (253.27±5.97°, n=174, circular mean±circular s.e.m) was not significantly different from that in RW_rich_ (p=1, circular Kuiper test). The mean vector length of the population (0.05) was not significantly different from chance (z=-0.34). **g,** The ensemble of head-directionally modulated neurons in 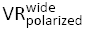, circular mean±circular s.e.m) were preferentially firing towards the visual cue (p=0.04, circular V test). Polarized Note the direction (111.27°) and the longer magnitude (0.12, z=2.50 which is significantly greater than chance) of the thick green line compared to (**e, f**). **h,** On the population level, neurons in 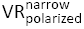, circular mean±circular s.e.m) were most biased towards the narrow cue (p=0.04, circular V test) as indicated by the magnitude (0.15, z=28.32) of the thick purple compared to all other conditions.

The absence of head-directional modulation in VR_blank_ may result from a lack of anchoring visual cues^32^ or optic flow created by the distal visual cues which could potentially be integrated to generate directional tuning. To address this, we performed another experiment where all the virtual walls had the same visual texture with high contrast and spatial frequency (VR_symmetric_), thus providing strong optic flow information but no angular information (Fig. 3c, see Methods). The virtual platform was placed in a larger room where each wall was 450cm away from the platform center, which ensured that the distance from the walls provided minimal spatial and angular information. Here too, neurons did not exhibit head-directional modulation (p=0.2, Wilcoxon signed-rank test), similar to VR_blank_ (Fig. 3d,i, Extended Data Fig. 5b). In fact, there was no significant difference between the degrees of angular modulation in VR_blank_ and VR_symmetric_ (p=0.6, Wilcoxon rank-sum test).

While internal mechanisms and optic flow may still modulate the degree of angular tuning, these experiments show that directional modulation is not generated by these mechanisms alone. This leaves open the possibility that head-directional modulation is generated by the angular information contained in the distal visual cues. To confirm this hypothesis we performed another experiment where the virtual world was strongly visually polarized. In this condition, there was just one high contrast wall, 450cm from the center of the platform, subtending a 90 degree angle 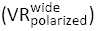(Fig. 3e, see Methods). This polarizing cue had no other spatial information and was identical to the walls used in the symmetric world. Here, 30% of hippocampal neurons showed robust head-directional modulation (Fig. 3f,i, Extended Data Fig. 6), which is a greater fraction than in all other conditions. However, the z-scored angular sparsity of neurons in 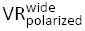 was comparable to that in RW_rich_ and VR_rich_ (Fig. 3j). Remarkably, the directional tuning curves of many neurons were much narrower (75.69±4.18°) than the sole, 90° wide polarizing cue.

Is there a lower bound on the width of the angular tuning curves? To address this we conducted another experiment where the sole polarizing cue was very narrow (11°), thus providing high angular information in the direction of the cue while leaving the majority of the maze blank 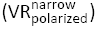(Fig. 3g, see Methods). Strong head-directional tuning was found in this condition as well (Fig. 3h, Extended Data Fig. 7). For neurons with significant head-directional tuning, the width of the tuning curves (70.94±5.34°) was not much narrower than in 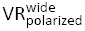 (Fig. 3k), and much wider than the 11° polarizing cue, indicating a lower bound on the width of hippocampal angular tuning curves. This could be influenced by internally generated hippocampal motifs enforcing a lower bound on the duration of the firing and hence the width of the tuning curves^11^. Further, the fraction of neurons showing significant head-directional modulation (19%) was considerably lower than in 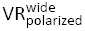 (Fig. 3i), perhaps because the narrow visual cue is visible to the rat for a smaller fraction of time than the wider polarizing cue, hence modulating a smaller percentage of neurons. Notably, despite similar z-scored angular sparsity for neurons with significant head-directional modulation in different conditions, z-scored mean vector length exhibited a different pattern which can be explained by the differences in the multimodality of the angular rate maps (Extended Data Fig. 8).

We then asked if the head-directional modulation of hippocampal neurons is stable and whether the stability depends on the experimental condition (Fig. 4). Tuning curves of neurons with significant head-directional modulation were significantly stable across the experimental session in all four conditions (RW_rich_ p=1.3×10^−43^, VR_rich_ p=2.0×10^−22^, 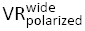 p=4.4×10^−16^ and 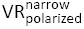 p=9.2×10^−12^, Wilcoxon signed-rank test)(Fig. 4a–d). The tuning curves were more stable (p=2.2×10^−9^) in RW_rich_ than in VR_rich_ (Fig. 4e,f). This could be due to the presence of other directionally informative multisensory cues in RW, such as distal odors and sounds, and their consistent pairing with visual cues resulting in higher stability. On the other hand, the tuning curves were more stable in the polarized VR experiments than in either of the rich conditions (Fig. 4e,f) indicating there may be competing influences of multiple cues within each modality in the rich conditions.

These results demonstrate that specific aspects of visual cues modulate the angular tuning of individual neurons; could they also influence the ensemble response? To address this we investigated the activity of the head-directionally modulated neurons on a population level under the four different conditions. For each neuron, the direction of maximum firing was computed from its angular rate map and was designated as its preferred direction (Fig. 5a–d, see Methods). We then computed the distribution of these preferred directions and the degree of angular bias of the population for each condition. There was no significant angular bias, as measured by z-scored mean vector length of the population (see Methods), in both RW_rich_ (z=0.95) and VR_rich_ (z=–0.34), and the two distributions were not significantly different from each other (p=1, circular Kuiper test) (Fig. 5e,f). The lack of population bias in the rich conditions is likely due to the presence of multiple visual cues on all walls, each contributing to tuning towards different directions.

Indeed, in 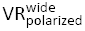 with only one visual cue, the population was significantly biased (z=2.50). This directional bias of the population was significantly (p=0.04, circular V test) oriented towards the prominent visual cue (Fig. 5g). The directional bias of the population was even stronger in 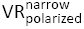, and was also significantly (p=0.04) oriented towards the narrow visual cue. There was an apparent reduction in the number of cells with preferred direction directly towards the narrow polarizing cue, and for some cells the preferred direction was opposite to the visual cue. This suggests that while visual cues drive the angular selectivity of hippocampal neurons, network mechanisms can modulate this activity.

## Discussion

These results demonstrate that, during two-dimensional random foraging, rodent hippocampal CA1 neurons show significant modulation as a function of head-direction with respect to the surrounding distal visual cues in both real and virtual worlds. This directional modulation does not require robust vestibular cues, while angularly informative visual cues are sufficient for its generation. Additionally, the directional modulation is strongly influenced by specific aspects of the distal visual cues, both at the neuronal and ensemble level.

Our demonstration of significant head-directional modulation of hippocampal neurons’ activity during random foraging in two dimensions in RW is contrary to the commonly held belief that head-directional modulation is absent in this condition^6,7^. The few reports about directionally modulated activity in hippocampus have reached conflicting conclusions, with some reporting vestibular-based head-directional modulation^3,14,25^ and others reporting vision-based spatial-view modulation^20–22^. By utilizing a VR setup we were able to isolate the contribution of only distal visual cues to hippocampal directional modulation. In addition, our analysis method was able to estimate the independent contribution of position and head-direction to hippocampal activity with high resolution, uninfluenced by behavioral biases^6^.

If hippocampal directional tuning was generated primarily from vestibular-based signals, one would expect a dramatic reduction of head-directional modulation in VR where the range of vestibular cues is minimized. However, hippocampal neurons not only showed significant levels of head-directional modulation in VR, but was also exhibited comparable levels to those in RW, thus demonstrating that robust vestibular cues are not necessary to generate directional tuning and that distal visual cues are sufficient.

About a quarter of all active CA1 neurons showed significant head-directional modulation in the visually rich RW and VR conditions, which is comparable to the fraction of directionally tuned neurons in several parts of the head-direction system^9^. We hypothesize that the directionally modulated hippocampal neurons may be the subset of the population that is predominantly driven by distal sensory cues, including distal visual cues. The rest could be largely driven by proximal cues, such as textures and odors on the track, which are not directionally informative. This hypothesis is consistent with prior studies showing a reduction in directional activity in the hippocampus with the inclusion of proximal cues^4^, and the remapping of subsets of hippocampal cells in accordance with the rotation of either distal or proximal cues^2,18,33,34^.

Head-directional modulation was abolished in VR in conditions with either no distal visual cues or symmetric distal visual cues demonstrating that it is not generated by internal mechanisms or optic flow alone. The directional modulation reappeared in VR with visually polarizing cues indicating that the angular information provided by visual cues directly influences head-directional modulation of individual hippocampal neurons. This influence was also observed at an ensemble level where the angular bias of the population was directly governed by the visual cues, i.e. no ensemble bias in the visually rich environments but a significant bias in the visually polarized cue conditions. Further, the degree of angular bias of the ensemble increased with increasing degree of angular polarization of the visual cue. Finally, this ensemble bias was pointed towards the polarizing visual cue. These results demonstrate a causal role of distal visual cues in driving hippocampal head-directional modulation.

Although a smaller fraction of neurons exhibited significant angular modulation in 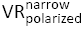 compared to the rich conditions, the degree of ensemble bias showed the opposite trend. We speculate that this is because, in the rich conditions, different neurons fire preferentially to different visual features on the walls, resulting in a greater fraction of angularly modulated cells and no ensemble bias. In contrast, in 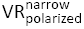 the visual features modulating neural responses were concentrated within a small range of angles, resulting in a reduction of fraction of directionally tuned neurons but a greater bias of the ensemble. These results indicate that visual cues do not provide a mere context for hippocampal activation, but rather that specific aspects of visual cues play a direct role in determining hippocampal responses.

Visual cues influence directional modulation of neurons in the head-direction system as well^9,35^. However, those neurons lose their direction selectivity without robust vestibular cues^36^, which is not the case in our hippocampal data from VR. Hence, we hypothesize that while the hippocampus may receive directional signals from the head-direction system, it must also be receiving directional information from a pathway that does not require the vestibular signal. This pathway could be through the parietal and retrosplenial cortices, which project to the hippocampus via the entorhinal cortex^9,37^ where neurons show significant head-directional modulation^38,39^.

In addition to the independence from vestibular cues, there are also other major differences between the properties of angular tuning curves in hippocampus and head-direction system. First, in hippocampus the tuning curves are much broader and more multimodal than those in the head-direction system^9,30^. Second, in the presence of a single polarizing visual cue, CA1 ensemble activity manifests a large bias in the direction of the visual cue, unlike the uniform distribution of preferred orientations in the head-direction system^31^.

Notably, the large reduction in spatial selectivity in VR^11^ is in contrast to the intact head-directional modulation observed here. This suggests that the mechanisms of spatial and directional selectivity can be dissociated, in that visual cues are sufficient to generate the latter but not the former. Further, these results bridge the gap between rodent, human and non-human primate studies showing the presence of angular selectivity independent of spatial selectivity^19,21,40,41^.

Our findings also narrow the gap between the presence of directionality on linear tracks^8^, but its apparent absence during random foraging in two dimensions^6,7^. Visual cues could generate the directional modulation of neurons in both one and two dimensions, and consistent pairing of visual and locomotion cues^11^ could enhance the degree of directional modulation on linear paths^3^.

These results could potentially resolve the apparent paradox: If the hippocampus is required for navigation^42^, how can rats navigate in a virtual world^10^ without hippocampal spatial selectivity^11^? We hypothesize that angular selectivity of hippocampal neurons, combined with their selectivity to distance traveled^5,11^, may be sufficient to solve a navigation task.

### Methods summary

Five adult male Long-Evans rats foraged for randomly scattered rewards in RW and various VR tasks. Four rats ran in visually similar RW and VR tasks with environments identical in size (300cm×300cm room with a 100cm radius elevated circular platform at the center). In addition, two rats ran on a 100cm radius platform in four other VR tasks with different distal visual features to determine their influence of hippocampal firing as follows: a) a room with no distal visual cues; b) a room with angularly symmetric cues with high spatial contrast positioned 450cm away from the center; c) a similar environment as in (b) but with only one high contrast cue subtending a visual angle of 90° at the center; d) a similar environment as in (c) but with the visual cue subtending only 11° angle at the center. Electrophysiological data were collected using bilateral hyperdrives with 22 tetrodes from dorsal CA1^5,11^. All procedures were in accordance with NIH approved protocols. Spatial and head-directional modulations were computed using a generalized linear model framework^15,27–29^. See online methods for details.

## Acknowledgements

We thank B. Popeney for help with behavioral training, J. Cushman and N. Agarwal for help with electrophysiology, P. Ravassard and A. Kees for technical support and help with surgeries, and B. Willers for discussions on analysis methods. This work was supported by grants to MRM from NIH 5R01MH092925-02, DARPA-BAA-14-08, and the W. M. Keck Foundation. Results presented in this manuscript were uploaded on a preprint server BioRxiv in March 2015.

## Online Methods

Materials and methods were similar to those formerly described^5,10,11^.

### Subjects

Data were obtained from five adult male Long-Evans rats (350–400g at the time of surgery) that were singly housed on a 12-hour light/dark cycle. Recording and training were both done during the light phase of the cycle. The animals were water restricted (minimum of 30mL/day) in order to increase motivation to perform the task, but allowed an unrestricted amount of sugar water reward during the task. Further, they were food restricted (minimum of 15g/day) to maintain a stable body-weight. All experimental procedures were approved by the UCLA Chancellor’s Animal Research Committee and in accordance with NIH approved protocols.

### Surgery, electrophysiology and spike sorting

All the methods were analogous to procedures described previously^5,11^. Rats with satisfactory behavioral performance were anesthetized using isoflurane and implanted with custom-made hyperdrives with 22 independently adjustable tetrodes. Both left and right dorsal CA1 were targeted. After recovery from surgery, tetrodes were gradually lowered to area CA1, which was identified by the presence of sharpwave ripple complexes. Signals were recorded using a Neuralynx data acquisition system at a sampling rate of 40kHz. Spike extraction, spike sorting and single unit classification were done offline using custom software and according to methods described previously^5^.

### Statistics

All analyses were done offline using custom codes in MATLAB. Two-sided nonparametric Wilcoxon rank-sum test and Kuiper test were utilized to assess the significance between linear variables and circular variables respectively. For tests of distributions being different from zero, Wilcoxon signed-rank test was used. CircStat toolbox^43^ was utilized to compute circular statistics. All values are expressed as mean±s.e.m unless stated otherwise.

### Random foraging tasks in visually rich RW and VR

These tasks were the same as those previously described^11^. Briefly, in both RW and VR, a 100cm radius elevated (50cm above the floor) platform was located at the center of a 300cm×300cm room whose walls had distinct visual cues as depicted in Fig. 1a (referred to as RW_rich_and VR_rich_). As commonly done, in RW rats foraged for food rewards scattered randomly on the platform. In a visually similar VR environment, rats foraged for randomly located but hidden reward zones. Upon entry into reward zones, a white dot (20–30cm radius) appeared and sugar water was dispensed through reward tubes. Data were collected from four rats in both RW_rich_and VR_rich_.

### Random foraging in VR tasks with visual cue manipulations

Two rats ran in four visually different VR environments, all of which had the same platform as described above. The reward zone was marked by a small (12.5cm radius) white disc only visible from a very short distance. Upon delivery of a reward, the reward zone moved to a new pseudorandom location on the platform. Visible reward zones were used to ensure uniform coverage on the platform and to avoid any behavioral biases that might be caused by the changes in visual cues.

In the first experiment, all distal visual cues, including the walls and the floor were eliminated (VR_blank_). The pattern on the platform was the only source of optic flow but provided no spatial or angular information (Fig. 3a).

In the second experiment, four identical and angularly symmetric distal visual cues, positioned along the four virtual walls of the square room, 450cm away from the center of the platform (VR_symmetric_) were added to VR_blank_. Despite high spatial contrast (to maximize optic flow) the distal visual cues did not provide any angular information due to the angular symmetry and high spatial frequency of the pattern (Fig. 3c). Further, the four walls were made infinitely tall to eliminate information about the corners. The large distance of the cues from the center of the table ensured that the cues did not provide any spatial information.

In the third experiment, only one of the high contrast visual cues used for VR_symmetric_ was placed 450cm from the center, creating a single wide polarizing cue which subtended a visual angle of 90 degrees at the center 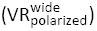 (Fig. 3e). This task had three variations where the visual cue appeared either in the front, right or left of the subject at the beginning of the session (Extended Data Fig. 6a,b,c). There was no quantitative difference between the data obtained in these variations, and hence these data were combined (data not shown).

In the fourth experiment, this visual cue was made narrower (11° visual angle) 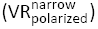 while maintaining the visual spatial frequency of its pattern, and placed at the same distance from the center as 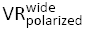 (Fig. 3g).

### Quantification of spatial and head-directional modulations using Generalized Linear Model

To quantify the influence of spatial and head-directional covariates on the firing of hippocampal neurons, and to minimize the influence of behavioral bias on spatial and angular selectivity estimates, we used a GLM framework^15,27–29^. The time-varying spiking activity was modelled as an inhomogeneous Poisson process as a function of space and head-direction:

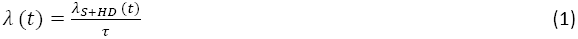

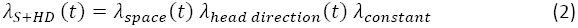

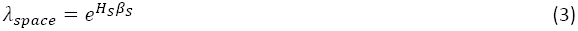

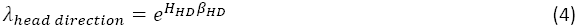

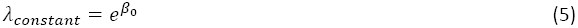

Where *τ* is the time bin size, *λ* is the intensity function, and *S* and *HD* denote space and head-direction respectively. *H*_*S*_ and *H*_*HD*_ refer to the design matrix associated with spatial and head-directional covariates and *β*_*S*_ and *β*_*HD*_ are the parameters associated with these matrices. Here *β*_0_ is a constant and the exponentiation is done element-wise. We expressed basis functions for *H*_*S*_ in terms of the set of orthogonal two-dimensional Zernike polynomials and *H*_*HD*_ in terms of sines and cosines. Equation 3 and 4 can be expressed as follows:

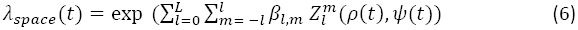

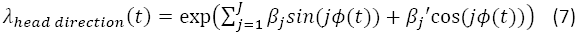

In equation 6, 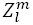 denotes the *m*^th^ component of the *l*^th^-order Zernike polynomial and *ρ*(*t*) and *ψ*(*t*) denote radial and angular components of position in polar coordinates. In equation 7, *ϕ*(*t*) is the head-direction of the animal. The parameters of the model (*β*_s_) were estimated using the GLM function in MATLAB to maximize the likelihood of the model. Further, we used Bayes Information Criterion (BIC) for model selection. The number of the basis functions used for equations 6 and 7 was chosen to minimize the following measure:

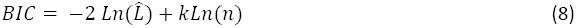

Where 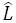 is the maximized value of the likelihood function of the model, *k* is the total number of parameters to be estimated across both space and head-direction, and *n* is the number of data points i.e. the length of intensity function. For a majority of cases BIC chose the fifth order in angle domain while this order was more variable in space domain. Hence, the number of angular basis functions was fixed at five. The number of spatial basis functions were allowed to vary and ranged from 5 to 32.

We then used the estimated parameters (*β*s) to reconstruct the modulation of the firing rate of neurons by spatial and head-directional covariates. For the reconstruction process and rendering purposes, we used 5×5cm spatial bins and a total of 80 angular bins (although the resulting rates are independent of these parameters). The reconstructed rates can be expressed as follows:

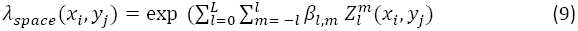

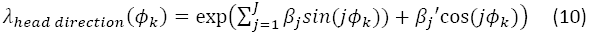

where *x*_*i*_, *y*_*j*_,refer to the spatial bins and *ø*_*k*_ refer to head-direction bins, and *β*_*S*_ are the estimated parameters from fitting. Thus, the spatial and angular modulation rates used in all the figures are defined as:

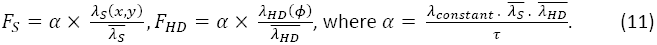

Here 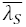 and 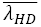 are the mean values of the spatial and head-directional reconstructed conditional intensity functions. To avoid artifacts, data from periods of immobility (running speed < 5 cm s^−1^) were discarded. During rate map reconstruction, bins with low occupancy time were excluded.

### Measures of selectivity

To quantify the degree of spatial and head-directional modulations we computed spatial and angular sparsity together with the mean vector length of the angular rate map. Sparsity of a rate map given *N*bins and *r*_*n*_ as the rate in the *n*^*th*^ bin is defined as:

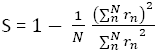

For firing rates as a function of head-direction, the mean vector length was computed as:

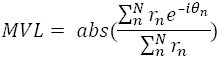

where *θ*_*n*_ and *r*_*n*_ are the angle and rate in the *n*^*th*^ circular bin respectively. Both of these measures are invariant to any constant scaling factor in the rates and hence remain unaffected by the normalization used when reconstructing rates using GLM framework.

## Generation of surrogate data to validate the GLM method

### Non-parametric generation of simulated place fields

To estimate the amount of angular modulation behavioral biases introduce into purely spatially modulated neurons, we generated surrogate data based on the firing rate maps of recorded neurons. Given a behavioral profile *B*(*t*) = (*B*_*X*_(*t*), *B*_*Y*_(*t*)) and spatial firing rate map *F*(*X*, *Y*), spike times were generated according to an inhomogeneous Poisson process with *F*(*B*(*t*)) as the rate parameter. Data generated in this manner were used in Fig. 1b and in Extended Data Fig. 1i.

### Parametric generation of simulated place fields

To verify the GLM framework accurately estimated the independent contribution of spatial and angular factors in determining spiking, we generated surrogate data with predetermined and variable degrees of spatial and angular modulation. For a surrogate place field centered at 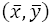, with spatial variance σ_*XY*_, preferred angular orientation 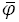 and angular variance σ_φ_, the relative probability of firing for any (*X*, *Y*, *φ*) combination was defined as:

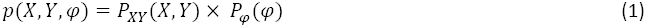

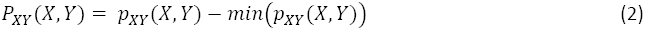

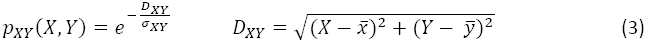

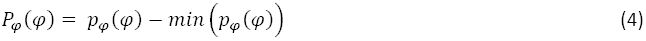

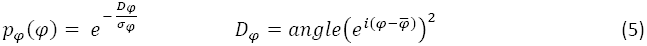

Where *i* is the imaginary number 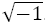.

Given a behavioral profile *B*(*t*) = (*B*_*X*_(*t*), *B*_*Y*_(*t*), *B*_*φ*_(*t*)) and desired mean firing rate *μ*, the absolute probability of firing is obtained by scaling the relative probability of firing (equation 1) by a constant factor k:

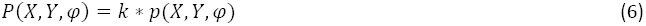

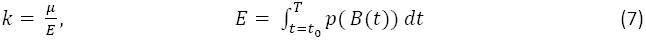

Where *t*_0_ indicates the start time of the session, and T indicates the end time of the session.

Spike times are then generated according to an inhomogeneous Poisson process with *P*(*B*(*t*)) as the rate parameter. Surrogate data generated in this manner were used in Extended Data Fig. 1a–h.

### Control analysis for spatial and head-directional modulations

To assess the statistical significance of neural modulation by position and head-direction, spike trains were circularly shifted with respect to behavioral data by random amounts (10–100s) to obtain control data. Spatial and head-directional modulations were quantified for the resulting rate maps, and for each neuron, all measures were expressed in the units of z-score or standard deviations around the mean value of this control data. Data exceeding two standard deviations were considered statistically significant at the 0.05 level. This method ensured that the degree of angular and spatial modulation was uninfluenced by nonspecific parameters such as the duration of the recording session and the firing rate of a neuron.

### Quantification of multimodality of angular rate maps

To detect the number of peaks in angular rate maps, a method similar to detection of place fields^5^ was used. First, all of the peaks with a minimum value of 50% of the global maximum were detected. For each peak, the boundaries were defined as the points at which rate drops below 50% of that peak for at least two angular bins.

### Quantification of significance levels of preferred firing direction of neural ensemble

For the head-directionally modulated neurons, preferred direction was defined as the direction of maximum firing obtained from the angular rate maps. To estimate the significance levels of the population bias, random angles between 0° and 360° were added to the preferred direction of the cells and the length of the mean vector was computed. This process was repeated 500 times and mean and standard deviation of these vector lengths were used to z-score the mean vector length in the experimental data. Z-score values greater than 2 were considered significant at the 0.05 level.

**Extended Data Figure 1:**
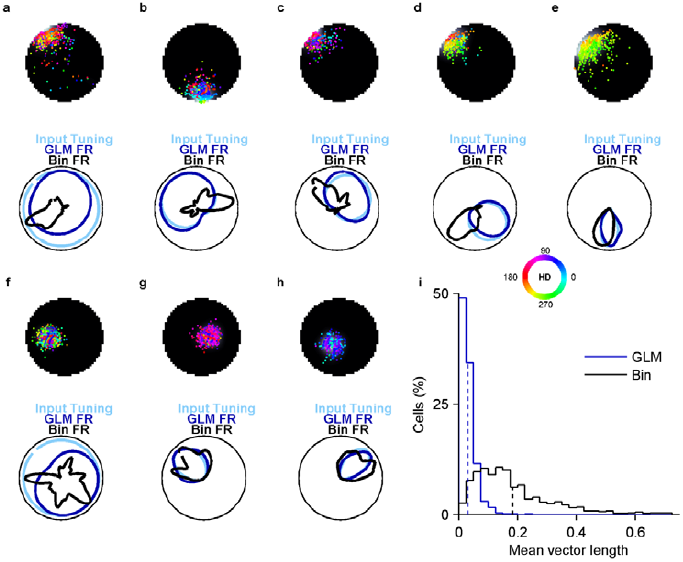
Comparisons of the performance of the GLM and binning methods in estimating head-directional modulation using surrogate data. Using experimentally recorded behavioral data, we simulated place fields by generating spike trains with arbitrary but adjustable spatial and angular modulations (see Methods). **a,** Top) Spatial firing rate (same color convention as in Fig. 1) of a simulated place field overlaid with colored dots representing the position at which spikes occurred. The color represents head-direction according to the color wheel (inset). Bottom) The angular part of the input function used for generating the simulated place field was uniform and had no head-directional modulation (light blue). The head-directional firing rate obtained by using the binning method (black) exhibited very sharp tuning, due to high behavioral bias at the edge of the platform resulting in a non-uniform sampling of the angles. In contrast, the GLM based rate (dark blue) followed the input function closely (showing no head-directional modulation). **b–h,** Same as in (**a**) but for other example cells simulated with different width and direction of input angular tuning. Note the similarity between the input tuning (light blue) and GLM based rate estimate (dark blue) in all cases, unaffected by the behavioral bias, which affects the binning method. **i,** Surrogate data were generated for each place cell with spatial modulation similar to that in experimental data but with no angular modulation (see Methods). Mean vector length obtained using the GLM method was close to zero (0.03±0.00, n=1066) and significantly (p=2.9×10^−278^) smaller (six-fold) than that computed using binning method (0.18 ±0.00). Thus, the commonly used binning method substantially overestimates the degree of angular modulation of spatially modulated cells, which the GLM method avoids. All values are reported as mean±s.e.m.

**Extended Data Figure 2:**
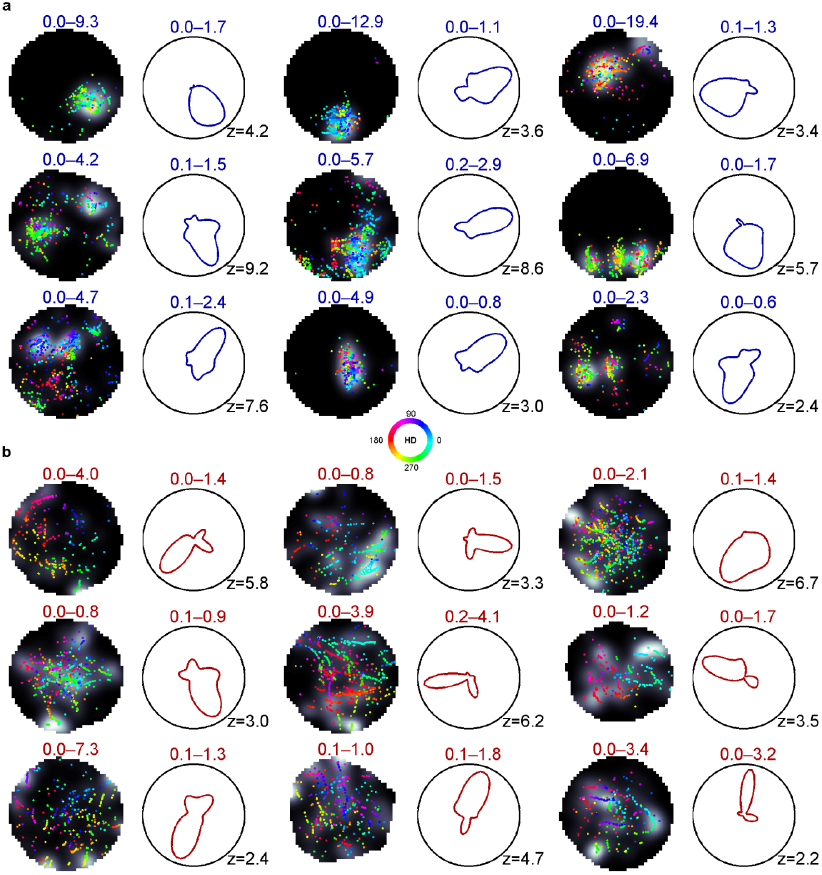
Sample cells in RW_rich_ and VR_rich_ with significant head-directional modulation. **a,** Spatial rate maps (grey scale, numbers indicate range) and spike positions (dots color-coded according to the head-directions) and head-directional firing rate (numbers in color indicate range, number at bottom right is z-scored angular sparsity of the angular rate map) of nine example cells in RW_rich_. **b,** Same as in (**a**) but for VR_rich_cells with significant head-directional modulation. Color conventions are the same as in Fig. 1 and will be used throughout the figures.

**Extended Data Figure 3:**
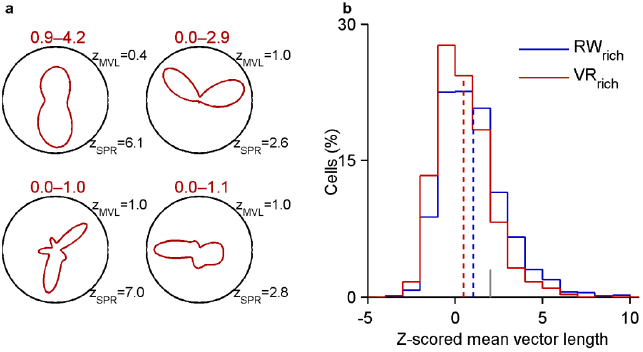
Unreliability of mean vector length in determining directional modulation of multimodal angular rate maps. **a,** Angular rate maps for four example cells in VR_rich_ with significant head-directional modulation determined from z-scored angular sparsity. All of them had smaller z-scored mean vector length due to multimodality of their angular rate maps (numbers in red indicate firing rate range; z_MVL_ and z_SPR_ correspond to z-scored mean vector length and z-scored sparsity computed from angular rate map respectively). **b,** This was reflected at the ensemble level where z-scored mean vector length of the angular rate maps in VR_rich_ (0.48±0.06, mean±s.e.m here and throughout, n=719) was significantly smaller (p=8.5×10^−10^, Wilcoxon rank-sum test) than cells in RW_rich_ (1.04±0.06, n=1066) despite identical z-scored angular sparsity (Fig. 2d).

**Extended Data Figure 4:**
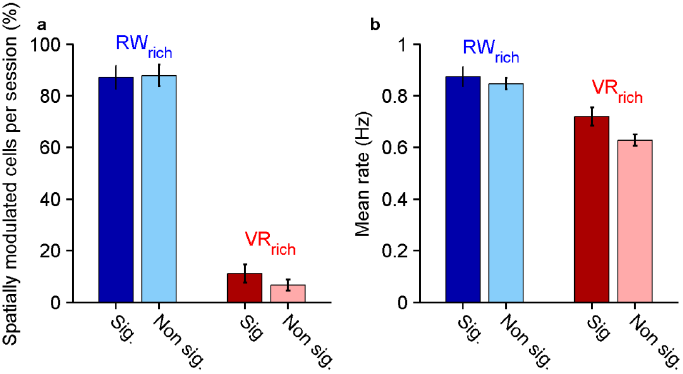
Comparison of spatial selectivity and mean activity of neurons with or without significant directional modulation in RW_rich_ and VR_rich_ conditions. **a,** In RW_rich_, the percentage of spatially modulated neurons per recording session was identical between neurons with or without significant head-directional modulation (87.12±4.49% and 87.87±4.13% respectively, p=0.6, Wilcoxon rank-sum test here and throughout figure legend). In VR_rich_, this percentage was slightly but not significantly higher for neurons with significant head-directional modulation (11.11±3.56%) compared to those which were not (6.67±2.19%, p=0.6). **b,** Mean firing rate of head-directionally modulated neurons in RW_rich_ (0.88±0.04 Hz, n=278) was similar to that in neurons with no significant modulation (0.85±0.03 Hz, n=788, p=0.3). In contrast, in VR_rich_, significantly head-directionally modulated neurons had higher mean rates (0.72±0.04 Hz, n=174) compared to neurons with no modulation (0.63 ± 0.02 Hz, n=545, p=8.7×10^−4^). Numbers are reported as mean±s.e.m and error bars indicate s.e.m.

**Extended Data Figure 5:**
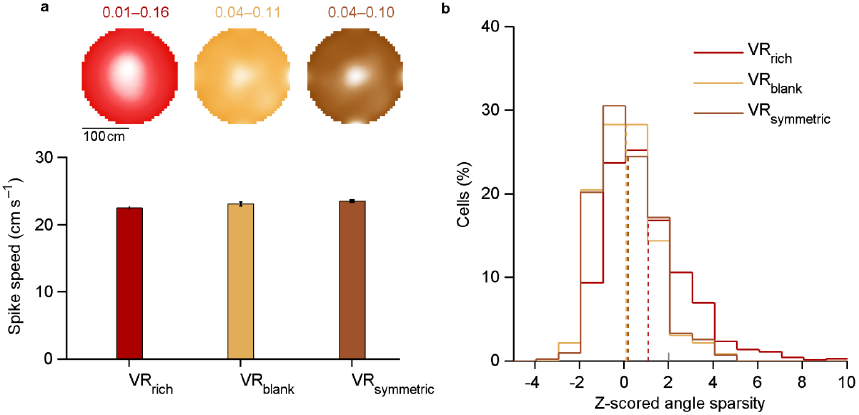
Similar behavior but absence of head-directional modulation in VR conditions with angularly uninformative cues. **a,** Top) Percentage of time spent in different parts of the platform in VR_rich_ (red), VR_blank_ (mustard) and VR_symmetric_ (brown). Numbers indicate range and lighter shades correspond to higher values. Bottom) Running speed in VR_blank_ (23.10 ± 0.30 cm s^−1^, n=230), was comparable to VR_rich_ (22.40 ± 0.13 cm s^−1^, n=719, p=0.03, Wilcoxon rank-sum test here and throughout this figure legend unless noted otherwise). Running speed in VR_symmetric_ (23.52 ± 0.17 cm s^−1^, n=426) was also similar but slightly greater compared to VR_rich_ (p=4.5×10^−7^). **b,** Z-scored angular sparsity of rate maps in VR_blank_ (0.09 ± 0.09, n=230) was comparable (p=0.6) to that in VR_symmetric_ (0.16 ± 0.06, n=426). Both were significantly less than VR_rich_ (1.09±0.08, p=1.3×10^−11^ VR_blank_ versus VR_rich_, p=1.4×10^−14^ VR_symmetric_ versus VR_rich_). While this distribution was significantly different from zero for VR_rich_ (p=3.0×10^−37^, Wilcoxon sign-rank test), it was not significantly different from zero in VR_blank_ (p=0.7) and VR_symmetric_ (p=0.2). Numbers are reported as mean±s.e.m.

**Extended Data Figure 6:**
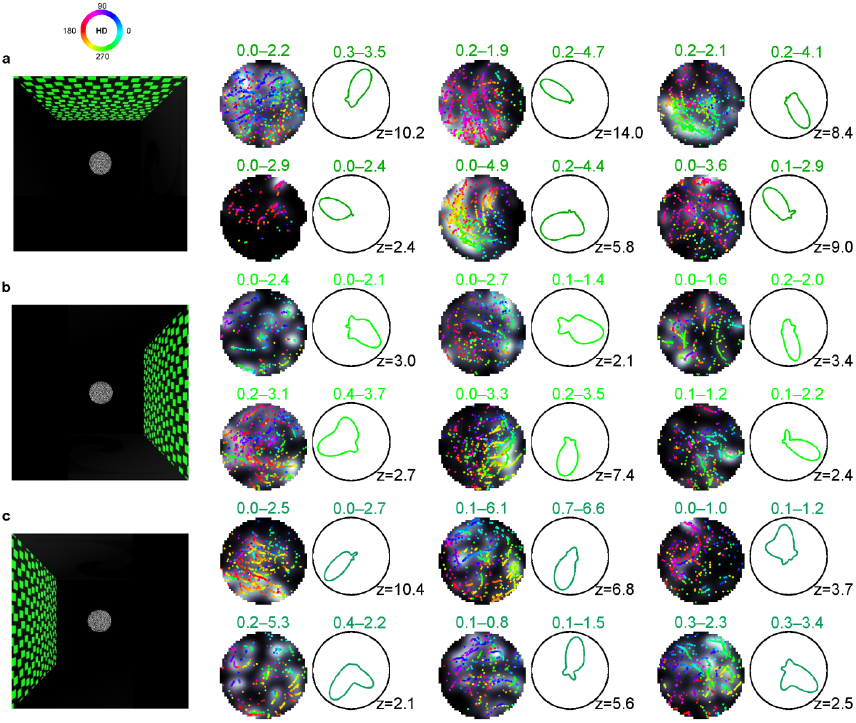
Example cells with significant head-directional modulation in 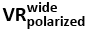 In different starting conditions. **a,** Left) Top view schematic of the virtual world with the polarizing cue located in front of the rat at the start of the session (color conventions as in Fig. 1). Right) spatial and angular rate maps of six different neurons (same conventions as in Fig. 1). **b,** Same as in (**a**) but for the task where the same polarizing cue is to the right of the rat at the start of the session. **c,** Same as in (**a–b**) but with visual cue located on the left side. All measures of head-directional modulation, namely z-scored angular sparsity and z-scored mean vector length were identical in these three starting conditions (p>0.5) and hence data from these experiments were combined. Direction selectivity is measured in the same fixed reference frame (color wheel at the top) in all conditions such that the polarizing visual cue is at 90°.

**Extended Data Figure 7::**
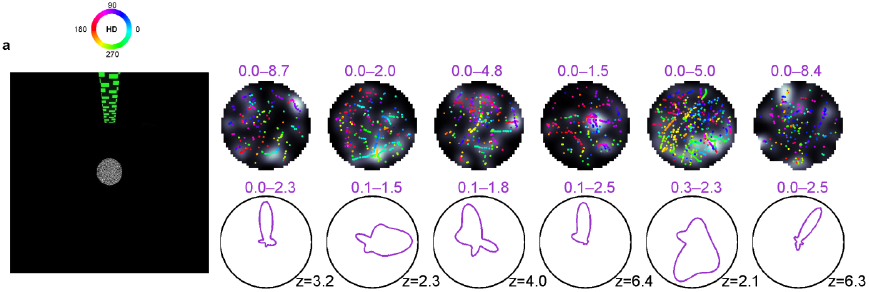
Sample head-directionally modulated neurons in 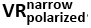. **a,** Left) A top-down schematic of the VR task with a narrow cue (11° visual angle) located 450 cm away from the center of the circular platform. Right) Spatial (top row) and angular (bottom row) firing rate maps of six different neurons (color conventions as in Fig. 1).

**Extended Data Figure 8:**
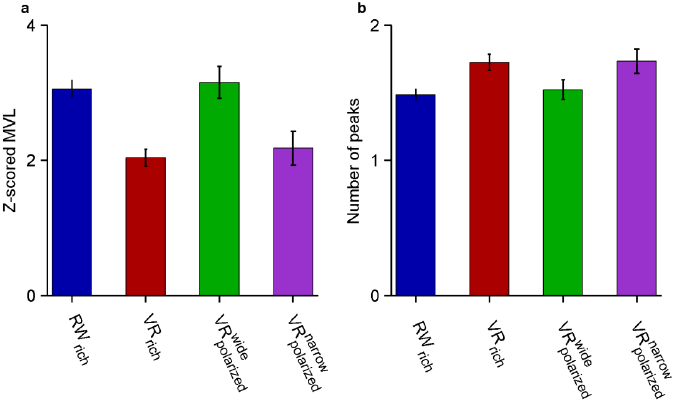
Quantification of mean vector length and multimodality of angular rate maps for different conditions. **a,** Mean vector length of the angular rate maps for head-directionally modulated cells in RW_rich_ (3.06±0.13), VR_rich_ (2.04±0.13), 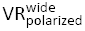 (3.15±0.23) and 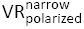 (2.18±0.25) are shown in the bar graph. For these, RW_rich_ mean vector length is significantly greater than VR_rich_ and 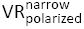 (p=2.0×10^−8^ and p=1.3×10^−3^ respectively). Similarly, mean vector length for 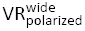 is significantly greater than VR_rich_ (p=4.7×10^−5^) and 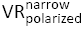 (p=6.9×10^−3^). Other comparisons were not significantly different from each other (p>0.5). **b,** Angular rate maps in VR_rich_ (1.72±0.06 peaks) were significantly more multimodal than RW_rich_ (1.49±0.04 peaks, p=1.5×10^−3^) and 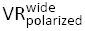 (1.52±0.07 peaks, p=0.04). The number of peaks in 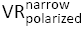 (1.73±0.05 peaks) was also significantly higher than RW_rich_ (p=8.2×10^−3^) and 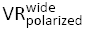 (p=0.04). Numbers are reported as mean±s.e.m and comparisons between distributions were done using Wilcoxon rank-sum test unless otherwise stated.

